# Women in Science and Technology: An Indian scenario

**DOI:** 10.1101/817668

**Authors:** Akanksha Swarup, Tuli Dey

**Affiliations:** Institute of Bioinformatics and Biotechnology, Savitribai Phule Pune University, Pune

**Keywords:** Women rights, higher education, gender inequality, women in STEM, India

## Abstract

The concept of treating women as equal to their male counterpart became a topic of political debate in Europe during the ‘age of enlightenment’ (18^th^ Century). The battle towards equal voting rights took approximately 100 years to win, and went until the 19^th^ Century. It was only around 1902-1920 that women got equal voting rights in prominent Western countries followed by others. Amidst the celebration of ‘women’s vote centenary’ throughout the world, the issue of equal rights to education and work is still waiting for the proper attention. Historically the presence of women in educational, technological and scientific fields remains mostly marginal. In this article, the current state of under-representation of women in the science and technology community is depicted, primarily highlighting the Indian scenario. It is observed that throughout India, and amongst the relatively developed countries of the world, the presence of women in highly prestigious Institutes and Universities remains negligible even in this day and age. The probable causes behind such inequality need to be analyzed, addressed and looked upon for remedial purposes.

## Introduction

With the emergence of ‘rational thinking’, ideas like ‘freedom from religion’, ‘abolition of slavery’, ‘constitutional government’ and ‘equal rights to women’ become central to the political debates during the “Age of Enlightenment”. The period also known as the “Century of Philosophy” starts loosely in 1620s with the scientific revolution in Europe and paved the way for different political revolutions of the 18^th^ and 19^th^ centuries ^1^. With the advent of 19^th^ century, New Zealand became the first country to grant voting rights to women followed by Canada, Great Britain and United States of America. India granted voting rights to women immediately after its freedom in 1947. The right to vote was quickly followed by the rights to equal employment and education, to provide better opportunities and social acceptance to women workforce. However, a brief glance of higher education and employment history shows very marginal participation of women, not just in India but also in the world. As a prompt example, the percentage of women Nobel laureates (2.94%) in the fields of Science and Technology along with the Field’s medal (1.66%) in mathematics gives us a brief idea. Such glaring inequity can be explained by many social prejudices and stereotypes towards women counterpart, questioning their intellectual and leadership ability. The age old concept of ‘women being the intellectually weaker section’ is often purported by many socioeconomical factors along with psychological stereotypes, such as.

a. **Gender stereotypes:** A stereotype of men being better at math and science is inculcated from childhood itself ^2, 3^. These preconceived notions might discourage female students to pursue their career in science or technology.
b. **Gender stereotyping of subjects:** There is also the longstanding belief that science and technology are masculine subjects, since they deals with the technical aspects of nature ^4, 5^. A recent study tried to explain the role of society and psychological impression over the masculinity of “physics” and thus highlighted the role of symbolic hegemony which perpetuated this idea ^6^.
c. **Social stereotypes:** Historically, the female intelligence was always believed to be inferior to male, fueled by Eugenics and Genetics ^7^. The hypothetical relationship between skull size and intellect supports the concept of intellectual inferiority of women ^2^. This school of thought persisted through time, and still does, despite several successful attempts to debunk it ^2, 8^.
d. **Role of women in family:** Another stereotype that plagues female worker in every sphere of life is the expectation to be the primary caregiver of the family.
e. **Lack of role model(s):** Lack of strong and assertive women characters, their negligible role in science and technology based popular culture media and literature might have played an important role in modulating the minds of girl children restraining them to follow such career paths.

Historically, the Darwinian concept that “the child, the female, and the senile white all had the intellect and nature of the grown up Negro” was profoundly fostered throughout the world ^9^. However, a brief analysis of the Indian history indicates that in early Vedic period (2000 B.C. to 1000 B.C.) women used to enjoy a prestigious status in science and education For example, Gargi Vachaknavi, ^10,11^ Lilavati ^12^, Maitreyi ^13^ were mentioned as the experts of their respective fields. However, such nurturing conditions did not prevail for long, with the onset of late Vedic and Epic period ^14^ (1000 B.C to 600 B.C). Additionally, ‘Arthashastra’ imposed even more stigmas on women during the Gupta period; further trimming away their already declining roles. The trend of women suppression continued during the Islamic and Mughal era, till late 18^th^ century to the mid-19th century, while India was mostly annexed by the British Empire. The prominent philosophers of western civilization such as Aristotle were found to advocate the inferiority of women ^15^. The documented history of Hypatia, the first female mathematician and astronomer of Alexandria, Egypt only survived till date ^16^. These early draconian concepts are further strengthened by Charles Darwin, in his treatise “The descent of man” and Francis Galton’s, “The Father of Eugenics”^17^. The World War I changed the scene for women in scientific research and a small, but substantial number of women joined the laboratories and engaged in science ^17^. India however, under the rule of the East India Company, observed many social reformers such as Raja Ram Mohan Roy, Ishwarchandara Vidyasagar, Jyotiba Phule and his wife Savitribai Phule, who struggled for the literacy of women. Inclusion of women in mainstream education and work force became more visible after the Independence struggle in 1947. The concept of women empowerment had been materialized in post independent India, where various social reforms were attempted for equipping them with rights to education and empowerment. However, it is observed that no such survey/data sets are available in the public domain providing the information about the ‘actual empowerment’ except few lone observations ^18^.

In the present study we tried to address that lack of information by looking into the current scenario of women in higher education pertaining to the fields of Science, Technology, Engineering and Mathematics (STEM) only. To understand the scenario, we have collected the information from various prestigious and prominent Indian Institutes/Universities and analyzed it. A comparative analysis of Indian Institutes of Technology (IITs) and the most prominent engineering colleges in the country depicts the immense gap in gender equality (Fig 1A) with an average female faculty of 11.24% (±4.65). The first group of IITs established between 1951-2001 (7 IITs) shows an average of 10.74% women faculty in science and technology streams, followed by 11.6% in second set (2008-2009) (8 IITs) and 11.33% in third set (2012-2016) (8 IITs). It can be assumed that the dearth of qualified female candidates remained the same from 1951 to 2018. However, the profile of faculty in the top twenty Universities of the country (Fig 1B) differed significantly as the average female faculty number increases to 27.11% (± 11.12). It can be probably explained by the fact that these Universities handle teaching and research mostly related to basic science rather than the technical subjects. Among the Universities, IISc, Bangalore came out as the mostly male dominated (with 8.6% female faculty), while Amritha Vishwa Vidyapeetham (45.8%) and Savitribai Phule Pune University (40.53%) stands out with most balanced distribution of gender within India. The National Institutes of Technologies (NITs) exhibit an average of 17.75% (± 5.65) of women faculty with NIT Uttrakhand having the lowest (6.67%) and NIT Manipur having the highest percentage (29.41%) of female faculty (Fig 2A). The Council of Scientific & Industrial Research (CSIR) is one of the largest research organization, with 38 national laboratories and approximately 4600 active scientists, distributed throughout India of which only 18.48%(±7.62) are women (Fig 2B). The distribution of gender in Indian Institute of Science Education and Research (IISERs) throughout the India is 15.47% (± 6.50) (female) and 84.52% (male) (Fig 3A) which is comparable to IITs and NITs. Different research institutes (mainly focused on biological sciences) such as NCBS, ILS, IICB etc, exhibit the same profile of having 23.12% (±9.74) female scientists compared to 77.49% (±8.88) of male scientists (Fig 3B). When studied, it became clear that irrespective of the year of establishment, eminence and geographical distribution, all the research and teaching institutes in India exhibited a similar pattern in employing female faculty/researcher (merely 10-20% of the total strength) with very few exceptions. Lastly, the distribution of female scientists/faculties amongst the top ten ranked Institutes throughout World showed an average participation of 20.34% (± 3.45) females which is marginally better than the Indian scenario (18.86%)(Fig 4A).

**Fig. 1.**
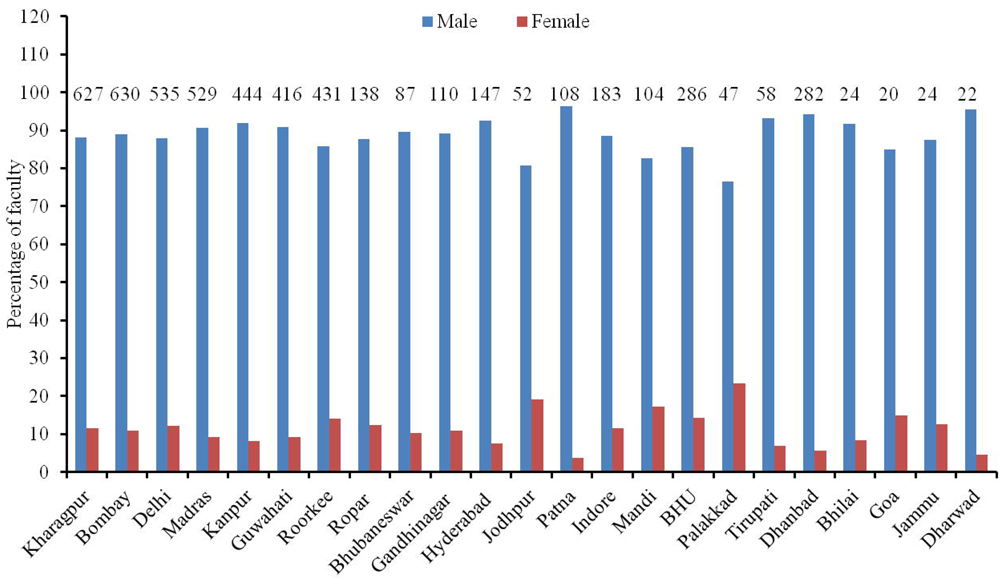
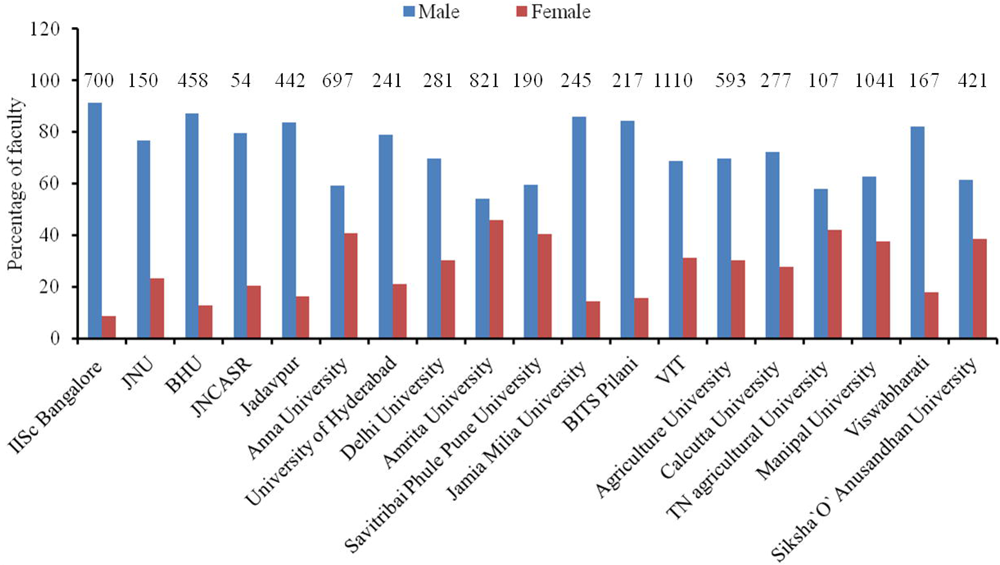
Comparative analysis of faculty distribution in IITs (A) and Top 10 Universities (B) in India. The Y axis values represent percentage of each gender and the numbers written on each bar exhibit the total number of faculties/scientists in respective Institutes.

**Fig. 2.**
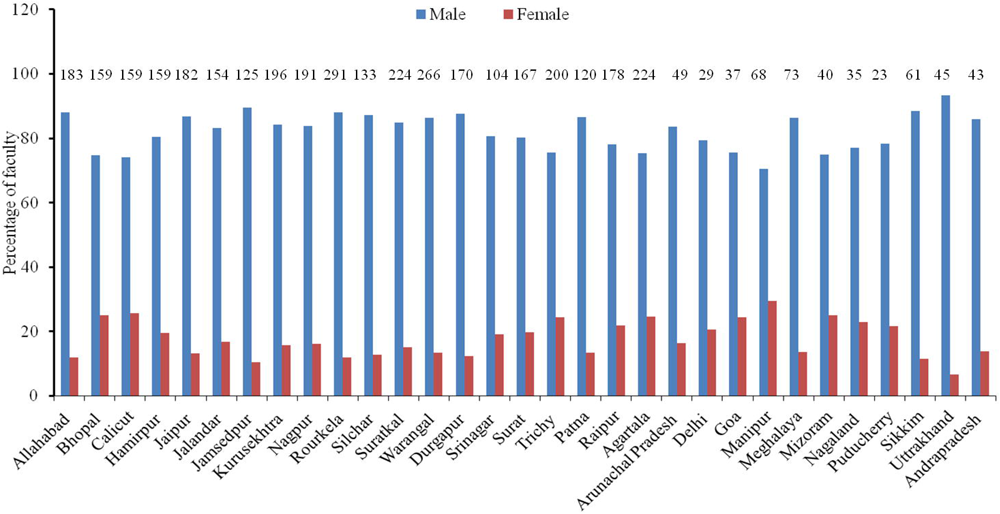
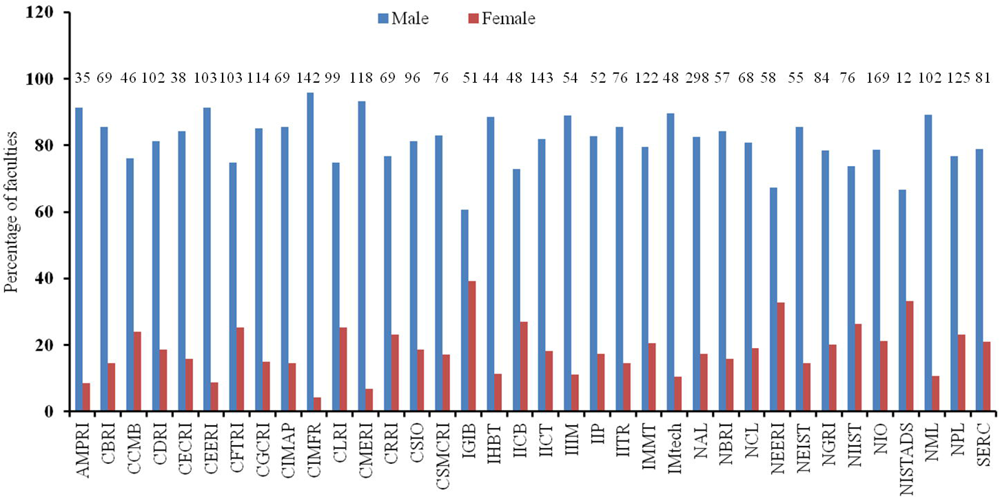
Faculty/Scientist distribution in NITs (A), CSIR Institutes (B) in India. The Y axis values represent percentage of each gender and the numbers written on each bar exhibit the total number of faculties/scientists in respective Institutes.

**Fig. 3.**
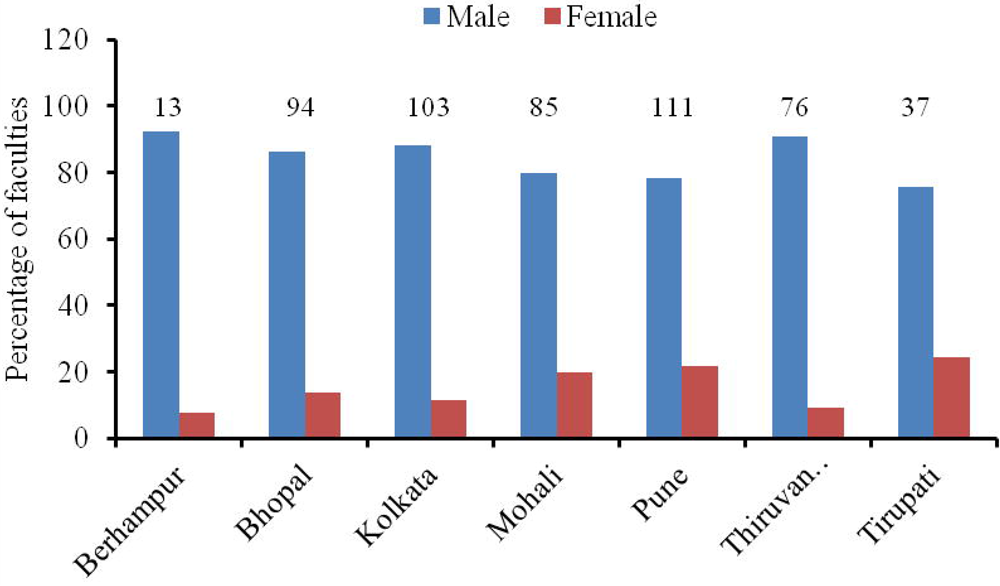
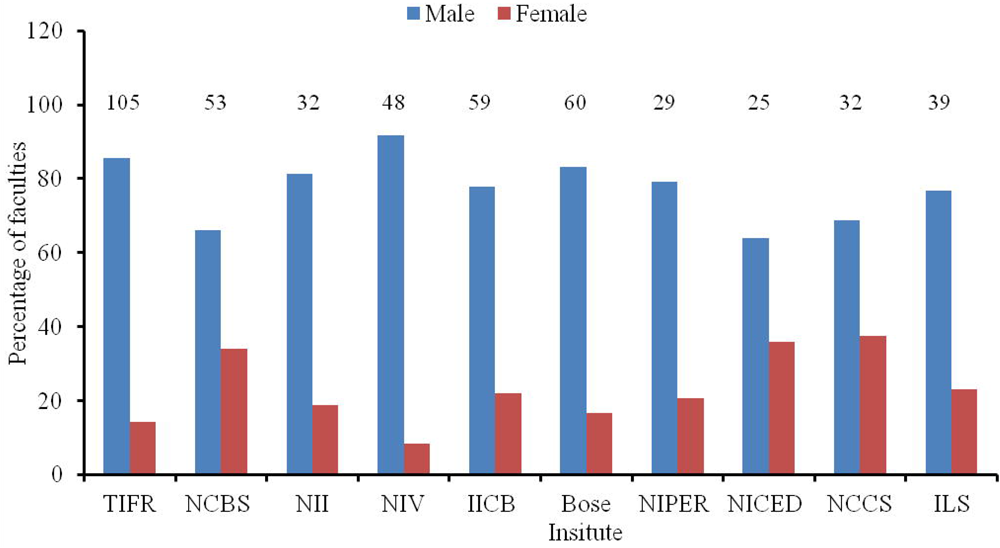
Gender wise distribution of total faculties in IISERs (A) and several significant biological science research institutes (B) in India. The Y axis values represent percentage of each gender and the numbers written on each bar exhibit the total number of faculties/scientists in respective Institutes.

**Fig. 4.**
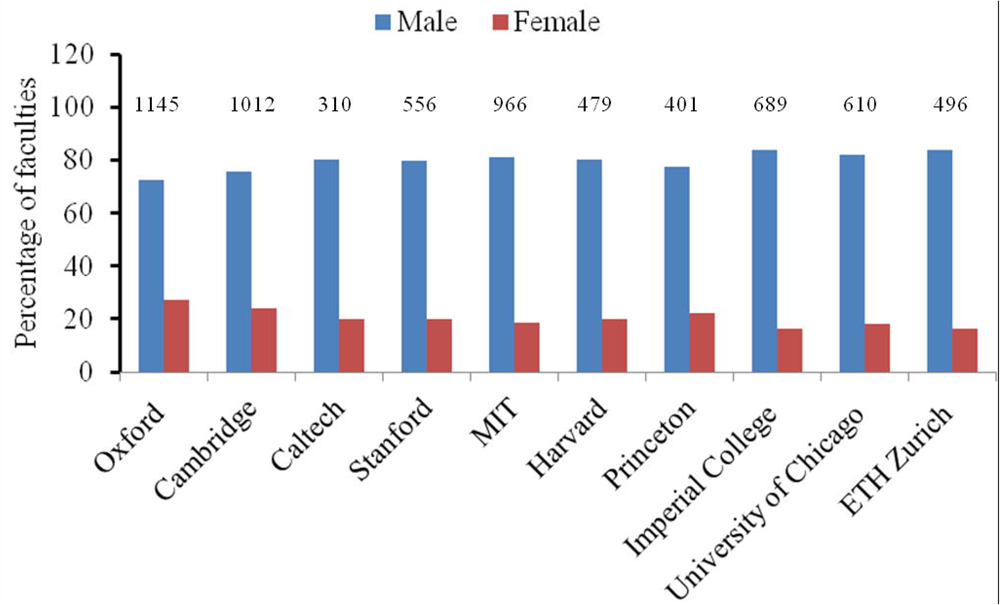
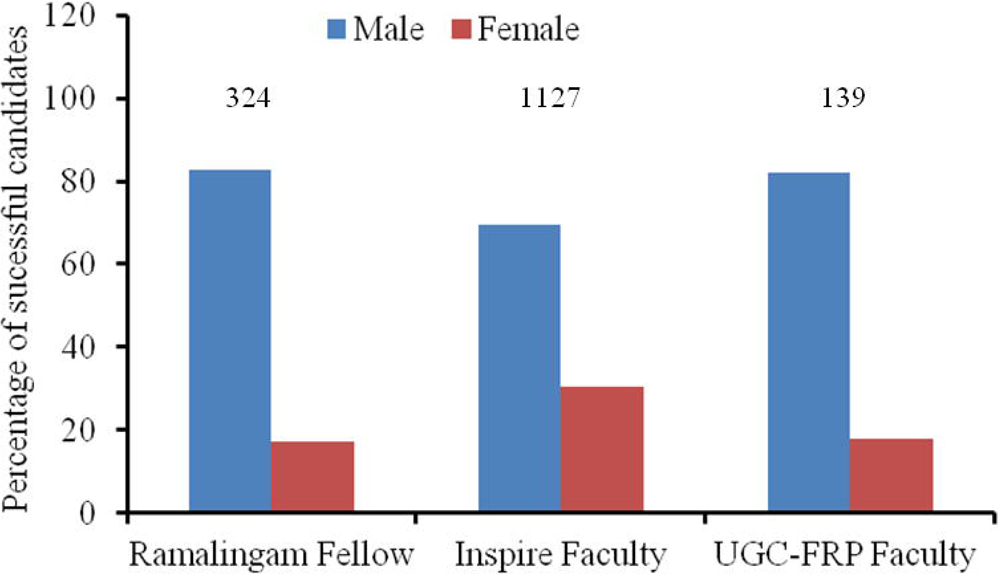
Comparative analysis of male and female faculties in top ten Institutes worldwide (A) and fellowship/faculty positions offered by different funding agency (B) in India. The Y axis values represent percentage of each gender and the numbers written on each bar exhibit the total number of faculties/scientists in respective Institutes.

**Fig. 5.**
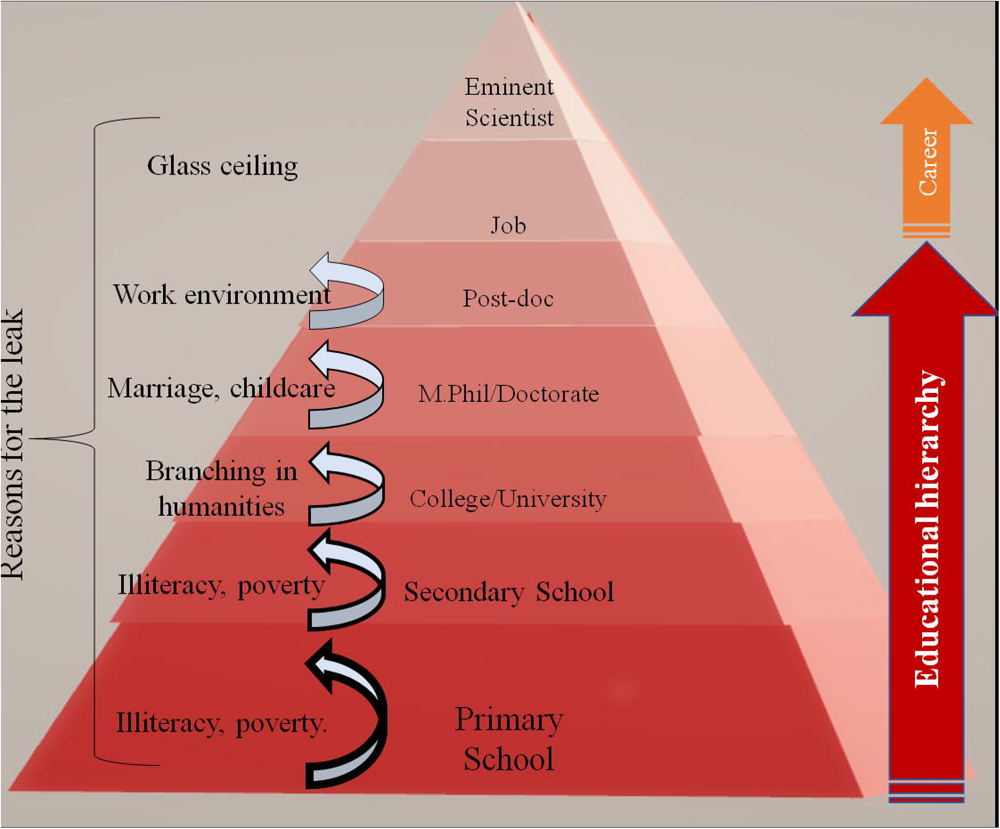
Schematic diagram of the concept of the ‘leaky pipeline’ in the education system. The right panel exhibits different stages of education, while the left panel denotes the probable causes of losing the girl students over time.

Such a huge gap between the male and female faculty distribution in the STEM cannot be reasoned only by not having enough female student to start with. The minimal number of female candidates pursuing a career in STEM can be explained by a model, called the “leaky pipeline effect’ ^19, 20^ where the leaking happens at primary school, secondary school, undergraduate, postgraduate, doctorate, post-doctorate and faculty level positions (Fig 5). The two of the greatest leaks in this system are transitions from secondary school to college and from post-doc to faculty positions due to familial and societal constraints ^20^. This phenomenon is further supported from the data (Fig 4B) where different research/Re-entry Fellowships/faculty positions, awarded in India over last 5 years, showed comparatively better female representation (21.88%) than the Institutes/Universities. Participation of women in STEM related jobs in the USA also shows minimal increment from 22% (1993) to 25% (2010) ^21^. It is, however important to note that there is no conscious decision in operation to filter out women in this system. However, the continuous loss of female workforce can be assumed to be a cumulative effect of the many factors ^2^ such as the followings.

a. **Social responsibility:** The absence of women both in primary and higher ranks of scientific community could be reasoned primarily by the family responsibility ^22^.
b. **Glass ceiling at work place:** Furthermore, due to these ingrained stereotypes supported by society and other scientists, women in science became a victim of self inflicted inferiority complex ^18^ and reduced presentation in senior authorship ^23^.
c. **Lack of recognition and Matilda effect**: The “Matilda effect” which refers to the prejudice against crediting women scientists for their work, and attributing the said work to their male colleagues might also have negative impacts ^24^. Multiple examples of this effect include renowned female scientists not duly credited for their ground breaking experiments such as Agnes Pockels, Nettie Stevens ^25, 26^, Frieda Robscit-Robbins ^27^ Rosalind Franklin ^28^, C.S. Wu, Jocelyn Bell, and Lise Meitner d) **Less participation in networking: It i**s observed that during scientific conferences, male researchers ask 1.8 times the number of questions asked by the same number of female researchers, which might impact the scientific networki^ng^ negatively ^29^.

### Probable solutions

General improvement in social and financial front especially in developing countries like India is paramount for more inclusion of women in main stream education which can be expected to trickle down further in STEM. Inclusion of women role model specifically from STEM should be more in popular media and scientific literature particularly in school textbooks might have positive impact. Scholarships, comfortable work place, flexible work hours, crèches and daycare facilities can be included to the current infrastructure to support the existing women workforce.

In spite of such hardships and obstacles, the scenario is changing worldwide, as 130% increase in enrollment of girl students was observed in male dominated subjects such as physics, technology, engineering, mathematics, statistics and computer science over the period 1990-2013 in the United States ^30^. In India, long term analysis is needed to understand the repercussions of previous policy changes and its impact on social structure.

## Methods

The data was collected primarily from the websites of the respective institutes, under the subpage titled “Faculty” and “Scientists”. Major institutes have been listed their faculty details under their respective Departments. The departments covered in this study were all STEM-related departments, such as physics, chemistry, mathematics and biological sciences (not including medicine). Departments not related to STEM, such as Management and humanities, while offered by the IITs and NITs (which give degrees focused on engineering) were not included in this study. The data was collected from the time period of May, 2018 to August, 2018. The numbers of awarded fellowships/re-entry fellowship/faculty positions by DST/Ramalingam/UGC were collected from the official sites within the period of Jan, 2019 to March 2019. Any change occurred there after would not be reflected in this study.

## Acknowledgements

The author would like to thank Dr. Vineeta Bal (IISER, Pune), Prof. Ameeta Ravi Kumar (IBB, SPPU, Pune) and Dr. Vaijayanti Tamhane (IBB, SPPU) for their for their critical reading of the manuscript and suggestions. The authors would like thank IBB and SPPU for infrastructure and support.

